# Hysteresis in calcium rhythm patterns during electric field stimulation of cardiac tissue

**DOI:** 10.1101/2020.02.29.970780

**Authors:** Harold Bien, Salmon Kalkhoran, Emilia Entcheva

## Abstract

Cardiac tissue subjected to fast pacing via electric field stimulation revealed hysteresis in calcium instability patterns (stimulus:response patterns) beyond departure and return to 1:1 response. The pacing frequency at which the first appearance of instabilities occurred (Fa) was higher than the frequency of ultimate disappearance (Fd) upon rate deceleration. Furthermore, hysteresis was observed in multiple pattern transitions. In the spatially extended system studied here, 2:2 alternans were the preferred starting point (Fa) in calcium instability development, while 2:1 blocks were more common in the return to Fa from higher pacing rates. Recovery of 1:1 patterns was preceded mostly by 2:2 alternans at Fd. In addition to previously reported hysteresis in action potential duration during 1:1 rhythm and alternans magnitude hysteresis (in 2:2 rhythm), our data reveal *hysteresis in rhythm pattern transitions* not just away from and return to 1:1, but also between different instability patterns, and thus provide insight into the rules of such transitions in electrically stimulated cardiac tissue.

## Introduction

Different stimulation history can prompt multivalued responses at a given pacing frequency during electric stimulation of cardiac tissue. This ambiguous frequency response hinders estimation of restitution properties for arrhythmia risk assessment or patient stratification and potential control attempts of life-threatening heart conditions (7).

*Hysteresis* in the response of cardiac tissue to electric stimulation is defined here as the appearance of a new pattern, e.g. alternans or irregular rhythms (**Alt/IR**), at a frequency **F_a_** during rate acceleration, and the disappearance of that pattern at a lower frequency (**F_d_** < **F_a_**) during rate deceleration. The difference between these two points has been often linked to short-term memory in the system (5;8;9). Hysteresis in response to pacing has been previously reported for action potential duration (APD) in 1:1 rhythm (9), for alternans magnitude in 2:2 rhythm (5;8), or for isolated patterns of destabilization, such as 1:1 to 2:1 transition (10). It has been attributed to slow time constants of recovery in the ion channel characteristics or to accumulation of calcium or other calcium-related events (8;11). The presence of hysteresis in patterns may indicate transient events (if steady-state is not established) or “true” *bistability* or *multistability*, where depending on the approaching stimulation path, different rhythm pattern may be observed in the **F_d_-F_a_** region (5;10).

In this report, we set out to examine the transition into **Alt/IR** region from flanking 1:1 and 2:1 rhythms during both rate acceleration and deceleration. Previously, we have extensively characterized intracellular calcium instabilities in cardiac tissue subjected to electric field stimulation within the **Alt/IR** region (1). Several classical pathways of destabilization, described in non-linear dynamics, were observed within that region. It was found that electric field stimulation of spatially-extended cardiac tissue is more likely to result in complex temporal dynamics (multiple rhythms) during pacing compared to space-clamped systems. Here we focus on the system behavior upon entry into and exit from the **Alt/IR** region. Thus, the goal was to characterize the observed patterns in the hysteresis region for cardiac tissue in hopes for better understanding of its frequency response.

## Methods

Cultured neonatal rat cardiomyocyte monolayers on rectangular strips 15×7mm (n = 35) were subjected to electric field stimulation (10V/cm, 2-3x threshold), **Figure 1**. Quiescent cells were paced starting from 1, 1.5, 2Hz and then in 0.2Hz steps towards and beyond the occurrence of alternans at frequency **F_a(up)_** and till occurrence of blocks **F_a2_**. This was followed by rate deceleration through **F_a(dn)_** till return to 1:1 rhythm (**F_d_**). Constant-BCL (basic cycle length) protocol was employed for pacing (80 beats per frequency), but only the last 20 beats after establishment of a pattern were used; “steady-state” was inferred by persistence of rhythm for all recorded beats without quantifiable monotonic change in the magnitude of the transients. Calcium transients were optically measured at 25°C with Fura-2AM at locations away from the electrodes (>3mm), and were classified as stimulus:response patterns as in (1).

**Figure 1.**
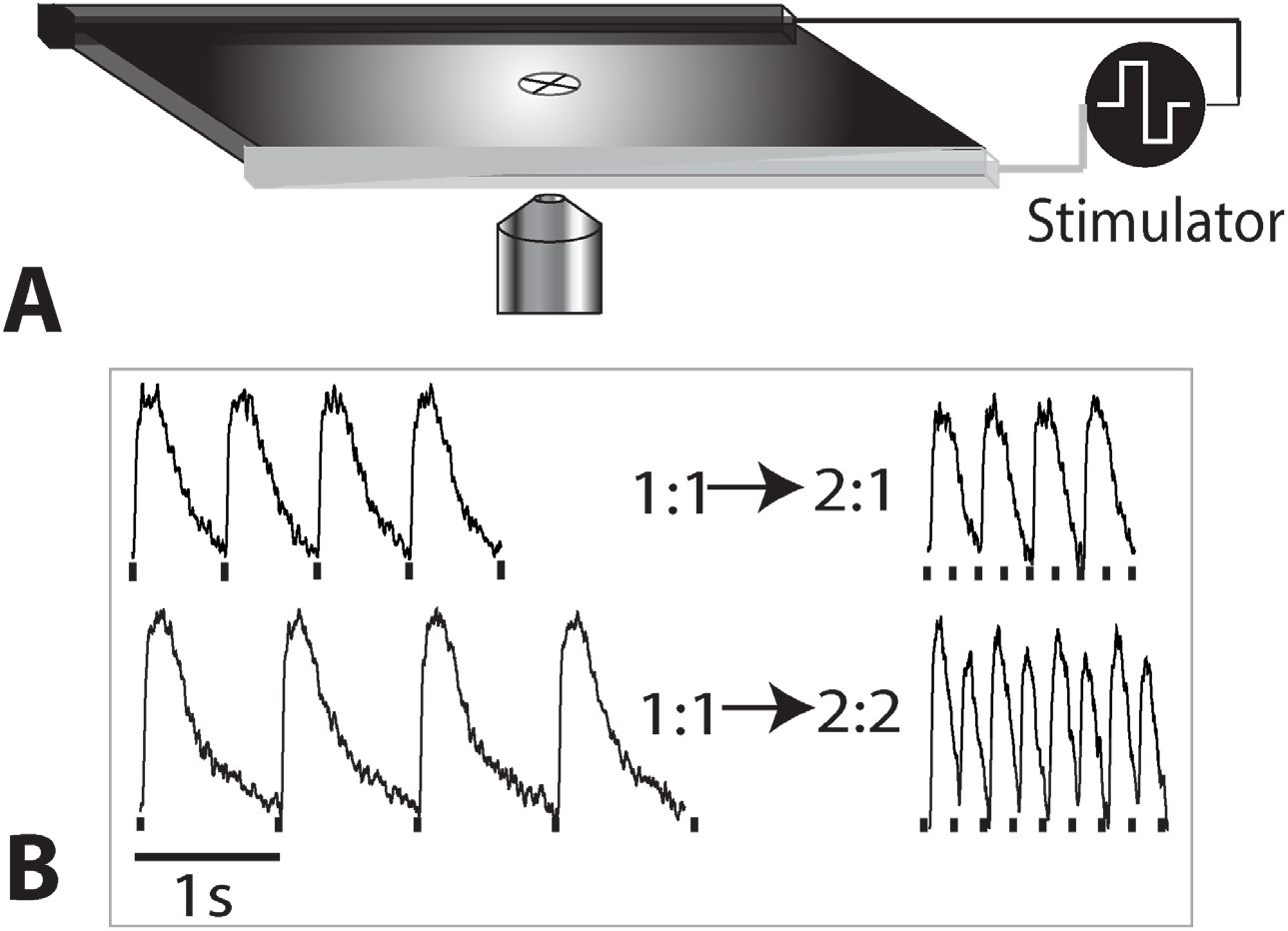
Schematics of the experimental setup (A), showing the sample, the filed-stimulating electrodes and the observation site. (B) Example traces from a sample undergoing direct 1:1 to 2:1 transition and from a sample undergoing 1:1 to 2:2 transition.

## Results

The most prevalent pattern at **F_a(up)_** during the transition from 1:1 to **Alt/IR** was 2:2 rhythm (52%), while upon return to **F_a(dn)_** the pattern typically has switched to 2:1 blocks (50%), **Figure 2**. Furthermore, samples returned to 1:1 rhythm at **F_d_** usually via re-occurrence of 2:2 alternans (63%). The majority of the other rhythms were Wenckebach N:(N-1) patterns. Multiple patterns were observed along with the 1:1 response in the **F_d_-F_a_** region, consistent with multistability if true steady-state is assumed at each pacing frequency. These data differ from previous studies with current injection in bullfrog tissue with measurements close to the stimulation site, where 76% of the patterns at **F_a(up)_** and 100% of the patterns at **F_d_** were 2:1 blocks (5). These differences could be explained by considering dissimilar restitution properties in the frog and the rat; taking into account coarser resolution in the frequency steps in previous studies, or considering the different measurements - voltage vs. calcium. Here, we propose overall left frequency shift in the rhythm patterns upon deceleration vs. acceleration and hysteresis in the 1:1 - 2:2 transitions, 1:1 – 2:1 transitions, as well as in the 2:2 - 2:1 transitions (**Figure 2**).

**Figure 2.**
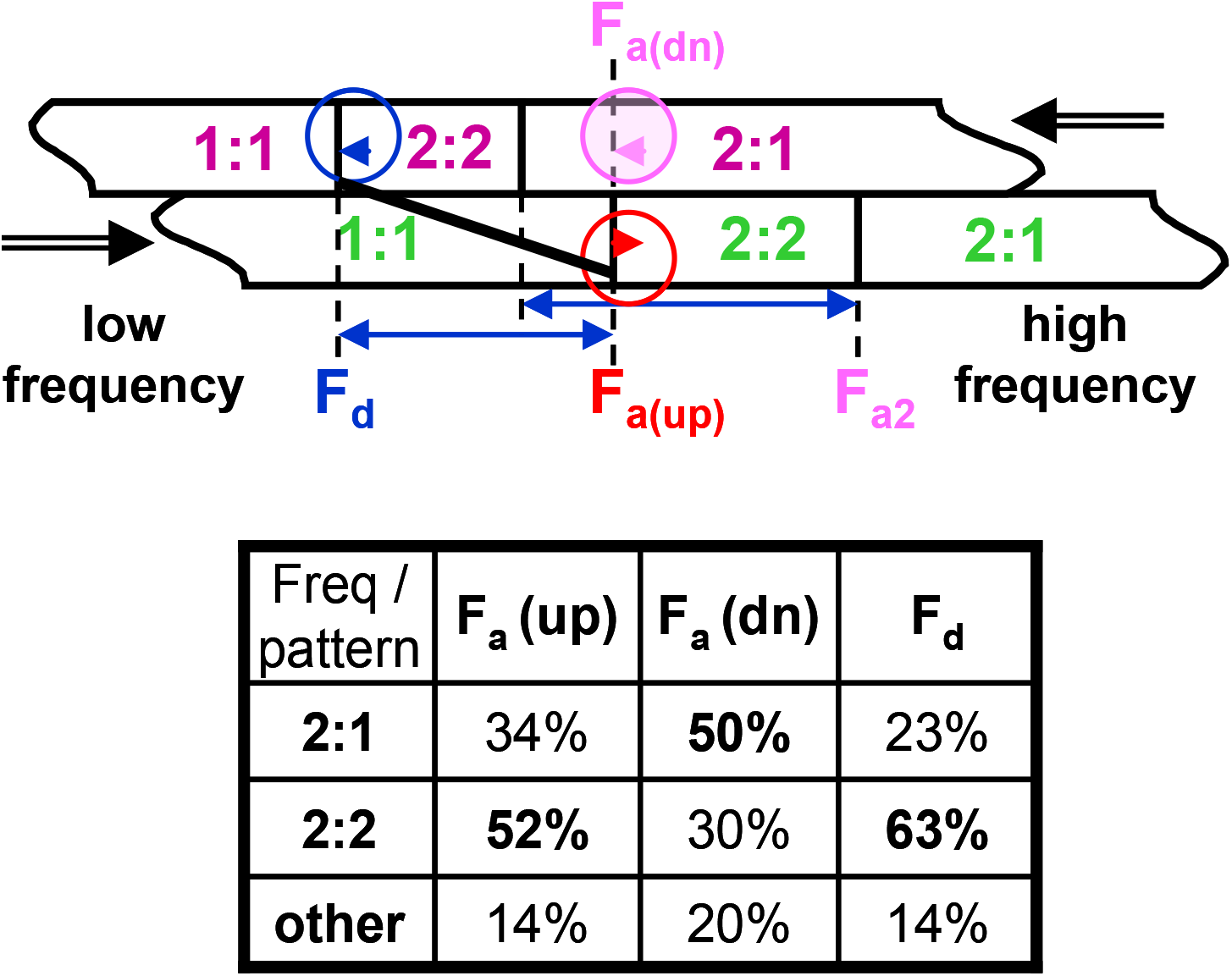
Proposed shift in patterns and observed prevalence during decrease vs. increase in frequency. Blue arrows indicate hysteresis in the 1:1 - 2:2 and in the 2:2 - 2:1 transitions. The table shows the prevalence of each pattern at the corresponding characteristic frequencies; the highest percentage is highlighted. In the “other” category, most common were Wenckebach patterns.

A staircase histogram of the behavior of a subset of 20 closely examined samples in **Figure 3** shows the proposed shift (hysteresis) in different pattern transitions over the examined frequency range 1 to 6Hz. First, upon rate acceleration, patterns transiently move from 1:1 to 2:2 rhythm and eventually to 2:1 blocks (**Figure 3A**). The bell-shaped 2:2 region reflects influx of samples from 1:1 and loss of samples to the 2:1 region. A left-shifted configuration similar to the one in **Figure 3A** will emerge upon rate deceleration. This frequency shift (hysteresis) is seen in each of the three curves (**Figure 3B-D**). The recovery of 1:1 rhythm, the persistence of 2:2 and 2:1 rhythms all extend towards lower frequencies compared to their frequency of occurrence, confirming *hysteresis in rhythm patterns*. The degree of 1:1 – 2:2 hysteresis in the examined samples was 29±6ms (mean±S.E.), while the 2:2 – 2:1 hysteresis was 18±3ms.

**Figure 3.**
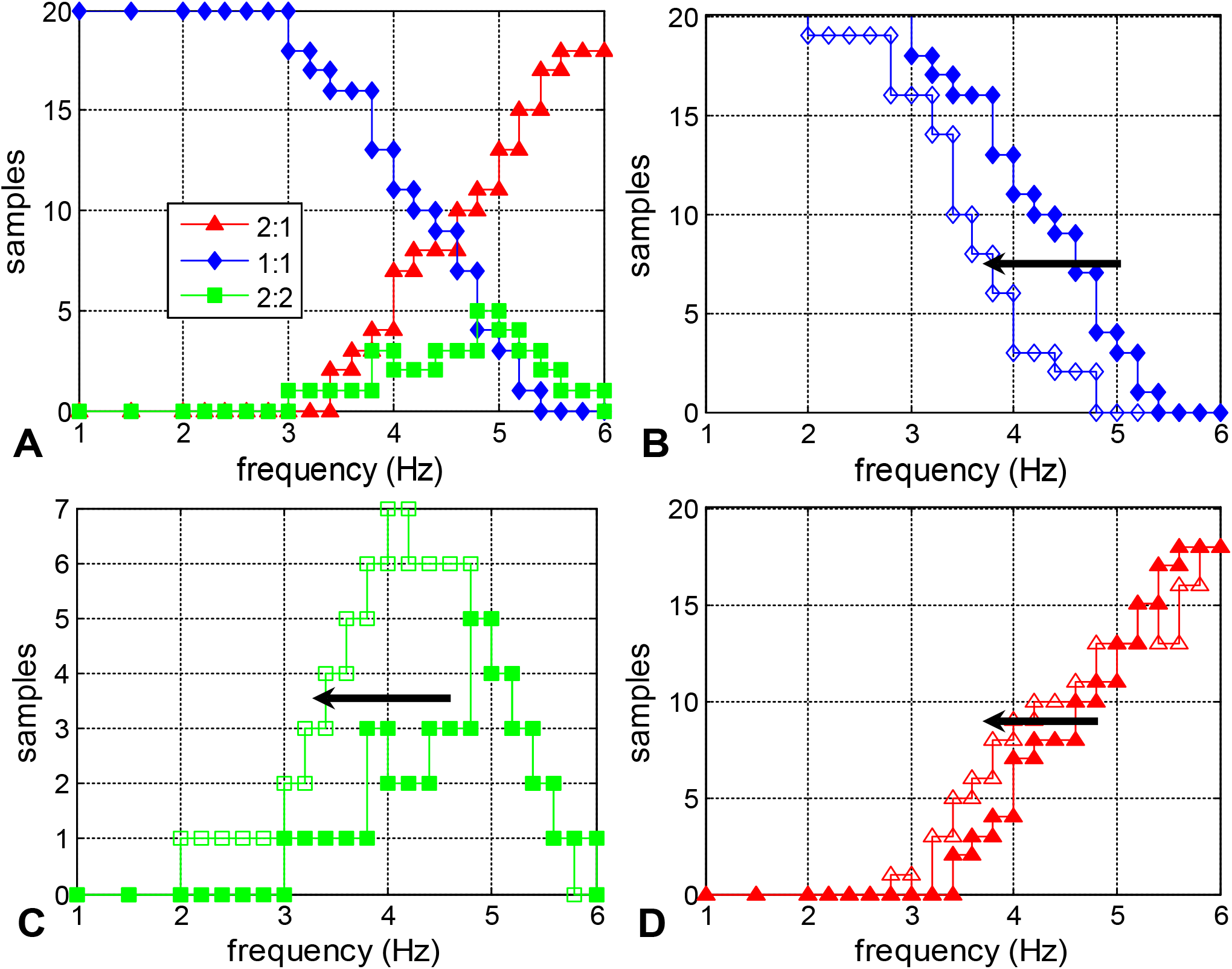
General distribution and hysteresis in patterns over a frequency range (1 to 6Hz). Staircase histogram plots in blue represent samples in 1:1 rhythm; red – samples exhibiting 2:1 block, and green – samples in alternans (2:2); filled and open markers indicate response to pacing with increasing or decreasing frequency, respectively. (A) Shown are only patterns in response to pacing with increasing frequency (up). Note the bell-shaped alternans region for intermediate frequencies during the switch from 1:1 to 2:1 rhythm. (B) Left frequency shift of the 1:1 rhythm curve for pacing down vs. pacing up. (C) Left frequency shift of the 2:2 rhythm curve for pacing down vs. pacing up. Overall rise in the 2:2 curve during pacing down is due to influx of samples from “other” rhythms, not represented in the three curves here. (D) Left frequency shift of the 2:1 rhythm curve for pacing down vs. pacing up. Black arrows indicate left frequency shift (hysteresis) in each curve.

It is interesting to uncover what dictates the choice of a particular path after loss of capture. In our system, for example, samples which experienced directly 2:1 block at **Fa(up)** (34%) had significantly longer calcium transients (*p*<0.01) in the low frequency range (1-1.5Hz) compared to those (52%) that underwent alternans (2:2), **Figure 4**. However, these differences disappeared at higher rates and the slope of the curves was not significantly different at the point of break from 1:1. Low frequency, periodic responses to electrical stimulation may hold clues to later dynamics.

**Figure 4.**
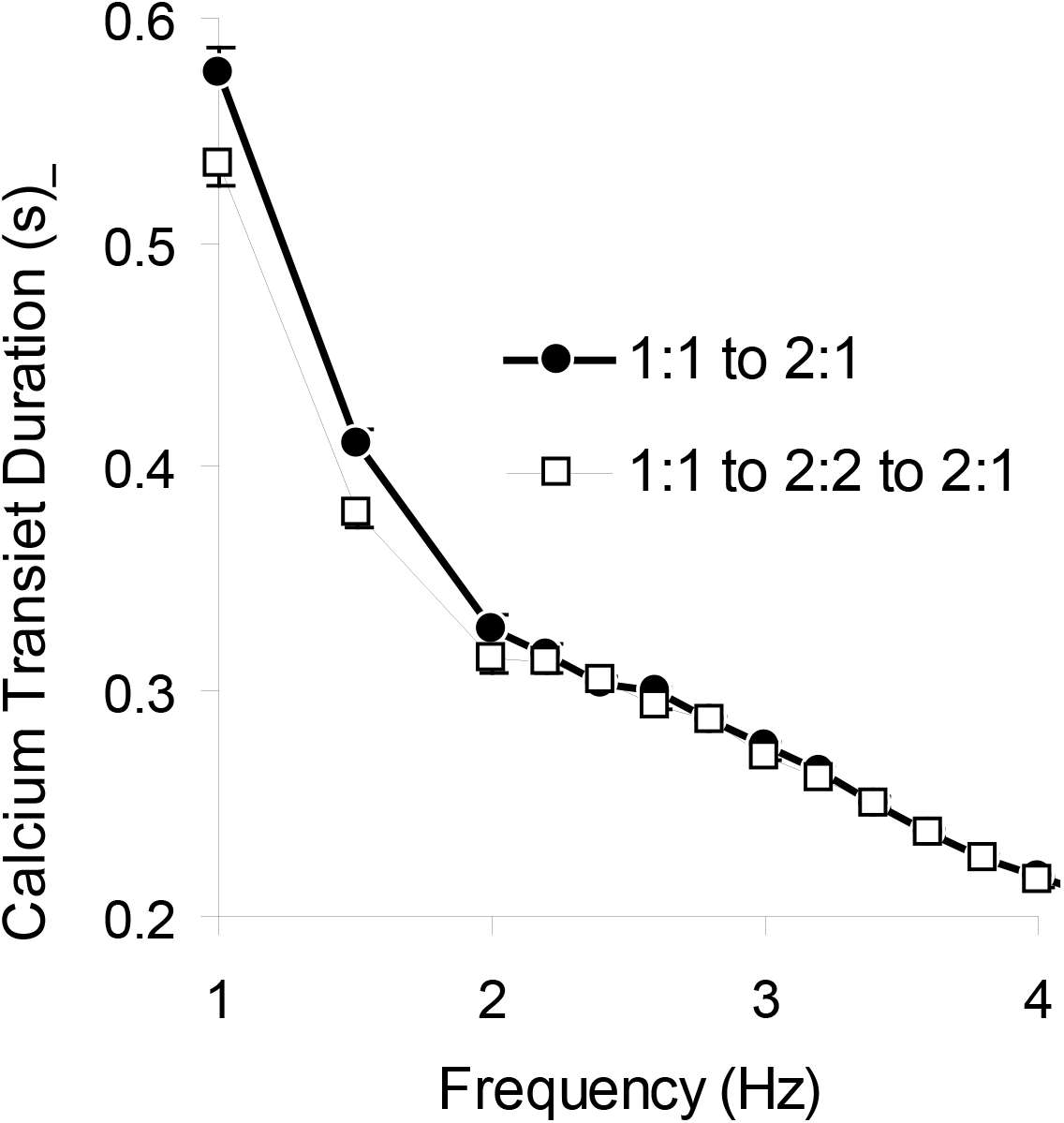
Frequency dependence of calcium transient duration (CTD) of samples falling into two categories: (a) those that directly went from 1:1 to 2:1 blocks at **Fa(up)**; and (b) those that followed 1:1 to 2:2 to 2:1 transition. Mean±S.E. are shown per frequency over all samples.

## Discussion and Conclusions

Our findings extend previously discussed hysteresis in APD during 1:1 response, hysteresis in alternans (2:2) or in isolated patterns (1:1 to 2:1) to a generalized view of hysteresis as a shift of patterns seen upon electrical stimulation of cardiac tissue on the way towards higher instabilities and back to 1:1 rhythm (**Figure 3**). This includes hysteresis in rhythm transitions other than departure from and return to 1:1, e.g. 2:2 - 2:1 transitions. Previously, theoretical work using one-dimensional maps obtained from APD restitution curves has predicted voltage hysteresis in these intermediate 2:2 – 2:1 transitions (4); furthermore instances have been reported for isolated myocytes (2). In addition, a study by Lorente (6) reported hysteresis in 1:0 – 1:1 transitions for sub-threshold stimulation. Nevertheless, our experimental study in a multicellular system is the first to date to examine and quantify hysteresis in the whole host of rhythm pattern transitions (1:1, 2:2, 2:1) in calcium within the same sample during supra-threshold stimulation.

When trying to understand the mechanism responsible for observations of hysteresis in cardiac tissue, one has to consider at least two possible scenarios. First, if the system is not allowed to reach steady-state within each pacing frequency step, it is possible that different transient rhythm patterns may be observed at the same frequency, but the system will eventually converge to the same stable state if left long enough at that frequency. In contrast, if “true” steady-state were guaranteed at each step, yet different rhythms are seen when increasing and decreasing pacing frequency, the system may exhibit bistability or “true” hysteresis.

In the cardiac literature, the duration required to achieve steady-state is not well known, nor characterized for our model system (or any experimental model, for that matter). However, in our experience, 80 beats are usually sufficient to establish a pattern, and offer feasible solution in fluorescence measurements used here (do not induce photobleaching during the course of the experiment). Our methodology is sensitive to gradual changes in alternans amplitude over time (20 recorded beats), and transient processes would have been visible had the system been trending towards a different state, e.g. 2:2->1:1 or 2:2->2:1. Since we believe that the observed behavior is not transient, we associate the hysteresis, reported here, with bistability, but cannot completely rule out transient response contributions, i.e. mixed response.

In the **F_d_-F_a_** frequency range, co-existence of alternans, hysteresis and Wenckebach rhythms was observed. This is in contrast to theoretical predictions for a space-clamped system, where strength of the stimulus delineates regions of Wenckebach rhythms (at low strength), hysteresis (at medium) and alternans (at high strength) (2–4). We believe that the variety of responses observed here reflects propagation phenomena and interactions in a spatially-extended system (1), where multiple effective stimulus strengths (and responses) can be experienced over space. Also, when interpreting our findings, one has to consider that calcium, not voltage, was measured and the dynamics of their interaction may not be trivial. In summary, our data support a generalized view of hysteresis in cardiac rhythm patterns during electric field stimulation, and may guide the definition of transition rules between such patterns for better understanding of life-threatening arrhythmias.

## Acknowledgements

This study was supported in part by grants from the National Science Foundation (BES-0503336) and the National Institutes of Health (R01HL144157) to EE.

